# Enhanced RNA Formation Under Amine-Rich Local Atmospheres from 2’,3’-Cyclic Nucleotides

**DOI:** 10.64898/2026.03.23.713775

**Authors:** Almuth Schmid, Aleš Kovařík, Jara Hintz, Sreekar Wunnava, Jan Palacký, Miroslav Krepl, Ondrej Šedo, Štěpán Havel, Katarzyna Ślepokura, Jiří Šponer, Peter Mojzeš, Christof B. Mast, Zbyněk Zdráhal, Dieter Braun, Judit E. Šponer

## Abstract

The core biopolymers (DNA, RNA and proteins) are assembled from their monomers under conditions that avoid water. RNA is crucial for the Origin of Life. When cleaved from its polymerized state, RNA first transitions to nucleoside 2’,3’-cyclic phosphates. In the reverse direction, RNA polymerizes from 2’,3’-cyclic monomers in dry states, creating short oligomers that then can ligate on a template under aqueous, alkaline conditions. We studied the role of the counterions in polymerization of 2’,3’-cyclic nucleotides under geologically plausible settings. Through experiments and simulations, we demonstrate that the presence of ammonium and alkylammonium counterions greatly improves RNA polymerization. The otherwise less reactive cytidine containing monomers formed polyC sequences of up to heptamers; copolymers of AU, GC, or GCAU were detected up to hexamers. Our findings suggest three reasons for this: (1) (Alkyl)ammonium cations form hydrogen bonds with phosphates, (2) their alkaline pK_a_ value can trigger general base catalysis, and (3) (alkyl)ammonium salts naturally form dry, anhydrous materials. The findings indicate that pyrolyzed organic tars creating ammonia-rich gas pockets in subsurface rocks could have enhanced the early evolution of RNA.

**TOC image:** 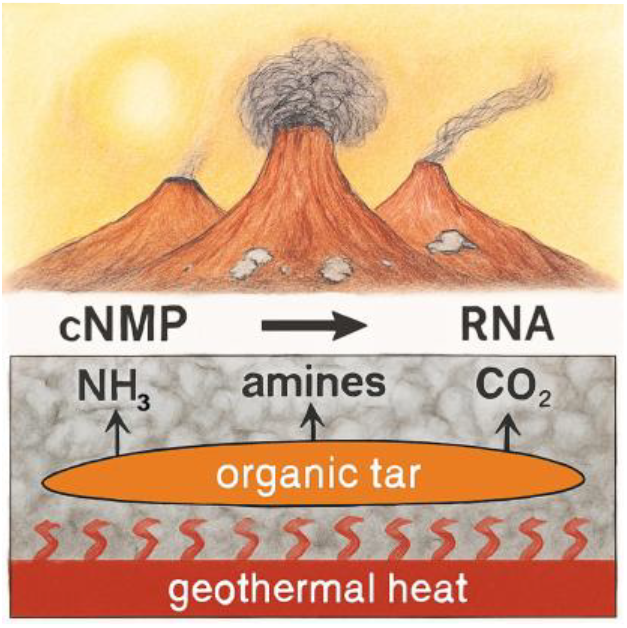

## Introduction

Nucleic acids, the fundamental constituents of living matter on Earth, form in condensation reactions from nucleotides by enzymes that very efficiently eliminate water from their active sites. RNA and DNA are essential for the storage and transfer of genetic information in all known forms of life. Understanding the mechanisms by which nucleotides can polymerize in the absence of enzymes and templates is crucial for elucidating the Origins of Life on Earth.

One of the main challenges in simulating prebiotic conditions is the removal of water,^1-3^ which is essential for promoting condensation reactions between nucleotides. Despite extensive research into various methods of nucleotide polymerization,^4-10^ these approaches often fall short due to the inherent hydrophilicity of the phosphate group in nucleotides. This hydrophilicity can significantly limit the efficiency of polymerization reactions, as it promotes degradation of activated nucleotide monomers.

Based on the available crystallographic reports^11-13^ it seems reasonable that the (alkyl)ammonium salt forms of nucleotides could offer a promising alternative to traditional metal salts to investigate template-free non-enzymatic polymerization of nucleotides in primordial conditions. Unlike the metal salt forms, which tend to incorporate structural water molecules into their crystals,^14^ (alkyl)ammonium salts are typically anhydrous in their dry forms. This property could facilitate higher polymerization yields and formation of longer oligomer chains in the absence of water.

In addition, the (alkyl)ammonium charge compensating cations could serve as general acid-base catalysts in the transphosphorylation reaction leading to internucleotide phosphodiester bond formation (see Figure 1).^10^ In the current work we focus on the polymerization of cytidine 2’,3’-cyclic monophosphate (cCMP), the least reactive member of the nucleotide series, to evaluate the effectiveness of (alkyl)ammonium salt form materials in promoting an acid-base catalyzed nucleotide polymerization. Besides elucidating the polymerization dynamics of cCMP, we gathered more insights into the potential pathways for the prebiotic synthesis of mixed oligonucleotide sequences containing all four canonical nucleobases.

**Figure 1.**
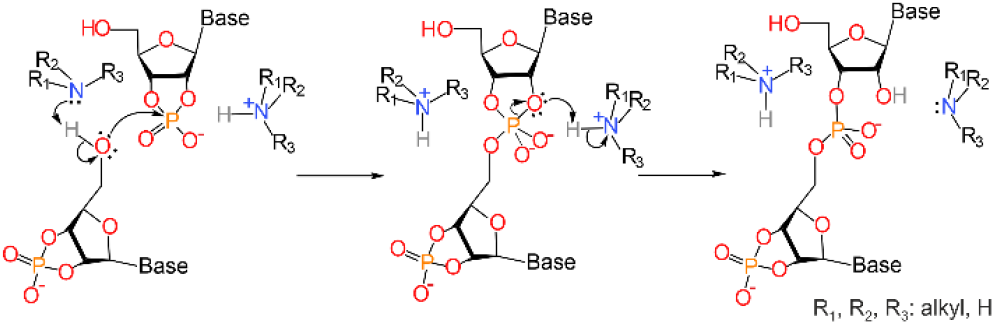
Mechanism. Proposed acid-base catalytic mechanism of the polymerization of 2’,3’-cyclic nucleotides with alkylamines.

## Results

### Comparison of the polymerization efficiencies of (alkyl)ammonium, sodium and potassium salts of cCMP in an ammonia/CO_2_/air mixture

The experimental setup used in the current study is shown in Figure 2A and is described in detail in the Experimental section of the Supporting Information. The polymerization reaction was conducted in a tightly closed Falcon tube containing a gaseous mixture of dry air, CO_2_ and NH_3_. The latter two gases were generated by thermal decomposition of ammonium carbamate, ammonium carbonate or other ammonium salt mixtures (*vide infra*). The nucleotide samples were dried on glass sample holders. The sample holders were immersed in the gas mixture that essentially served as a volatile NH_3_/CO_2_ buffer with a pH equal to that of ammonium-carbonate/carbamate, i.e. 9.25 (see Figure S1).

**Figure 2.**
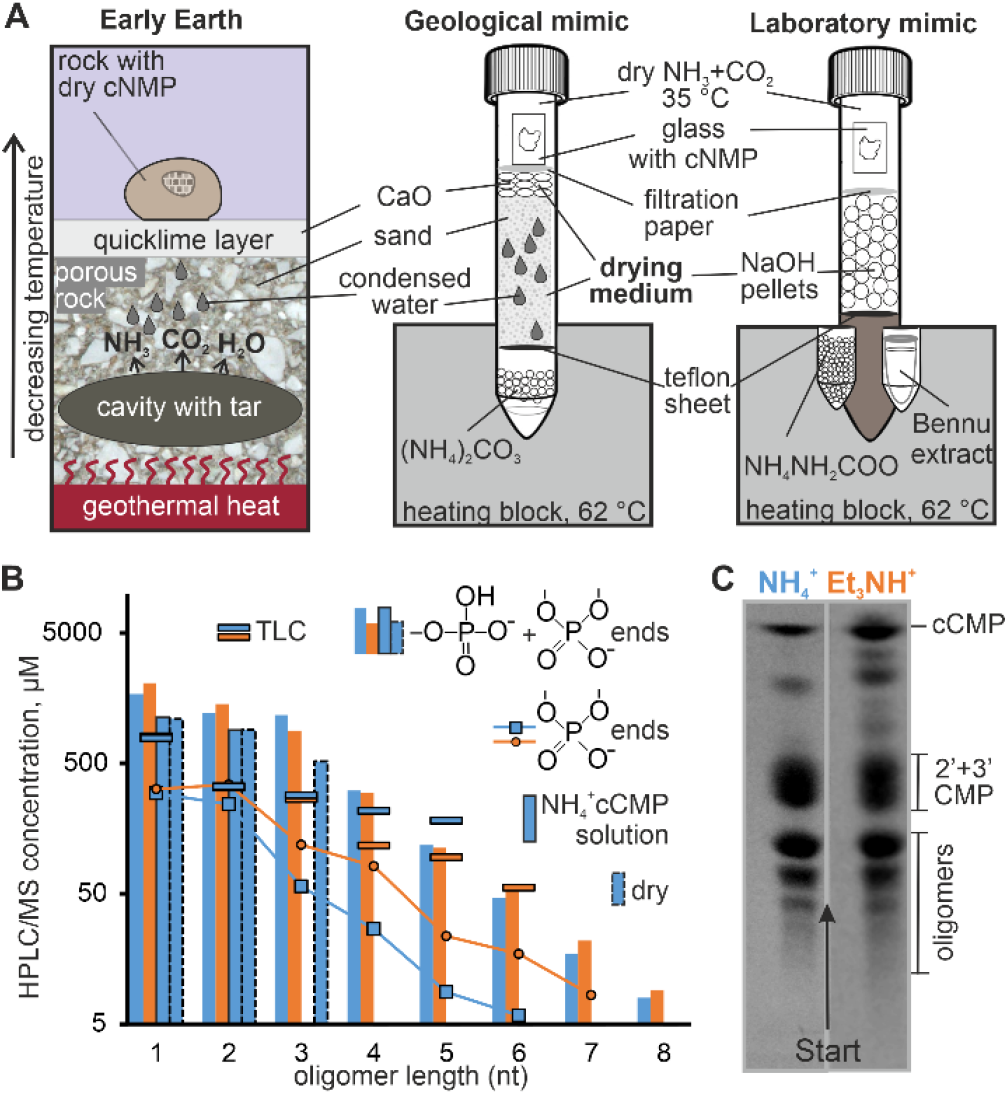
Counterions from amine-rich atmospheres enhance RNA polymerization. Polymerization of ammonium (NH_4_^+^) and triethylammonium (Et_3_NH^+^) forms of cCMP in an atmosphere containing ammonia or amines. A) Two different laboratory setups (referred to as “laboratory mimic” and “geological mimic”) were used to mimic a geological setting on early Earth shown on the left (“Early Earth”). In this geological model, a gas mixture of NH_3_ and CO_2_, formed by the pyrolysis of an organic tar, is released into the atmosphere through diffusion across porous rocks covered with a CaO (quicklime) layer precipitated on rock surfaces. The porous rock and the quicklime layer serve as drying agents to remove water from the gas mixture. In the “geological mimic” setup a combination of a thick sand layer and a thinner layer of quicklime was used as a surrogate of the drying agent. In the simpler “laboratory mimic” setup NaOH pellets were used for the same purpose. B) HPLC-MS length distribution of the products obtained by the polymerization of NH_4_^+^ (data in blue) and Et_3_NH^+^ (data in orange) forms of cCMP triggered by incubation in a vapor formed above ammonium carbamate using the setup “laboratory mimic” for 2 days. C) Thin-layer chromatographic analysis of a parallel sample used for the HPLC/MS analysis shown in panel B (for MALDI-ToF MS analysis of the materials isolated from the chromatographic bands see Figure S2 in the Supporting Information).

As Figures 2B-C and S2 (see Supporting Information) illustrate, when using ammonium carbamate to generate a gas mixture composed of CO_2_, NH_3_ and air, we detected oligomers in the length range of up to heptamers. About 80% of the cCMP monomers is incorporated in oligomeric form after a 2 days long sample treatment at 35 °C based on the peak intensities measured by HPLC/MS. Notably, though the decline of the polymerization reaction is still exponential with increasing oligomer length, it is remarkably slower compared to those found in previous reports on 2’,3’-cyclic nucleotide^10,15^ and imidazole-activated nucleotide^7^ polymerizations. At the same time, for 12% and 21% of the oligomers (for NH_4_^+^ and Et_3_NH^+^ forms of cCMP, respectively) the cyclic phosphate end is recovered after the reaction, leaving it with the possibility to further polymerize.

Figures 3 and S3 (see Supporting Information) compare the polymerization profiles of Na^+^, K^+^, and NH_4_^+^ salts of cCMP, as well as that of a 1:1 mixture of Na^+^ and NH_4_^+^ salt forms of cCMP. The data shows that the lowest polymerization efficiency is observed for the Na^+^ and K^+^ forms, and the amount of monomers incorporated into oligomers increases with increasing ratio of the NH_4_^+^ form in the samples.

**Figure 3.**
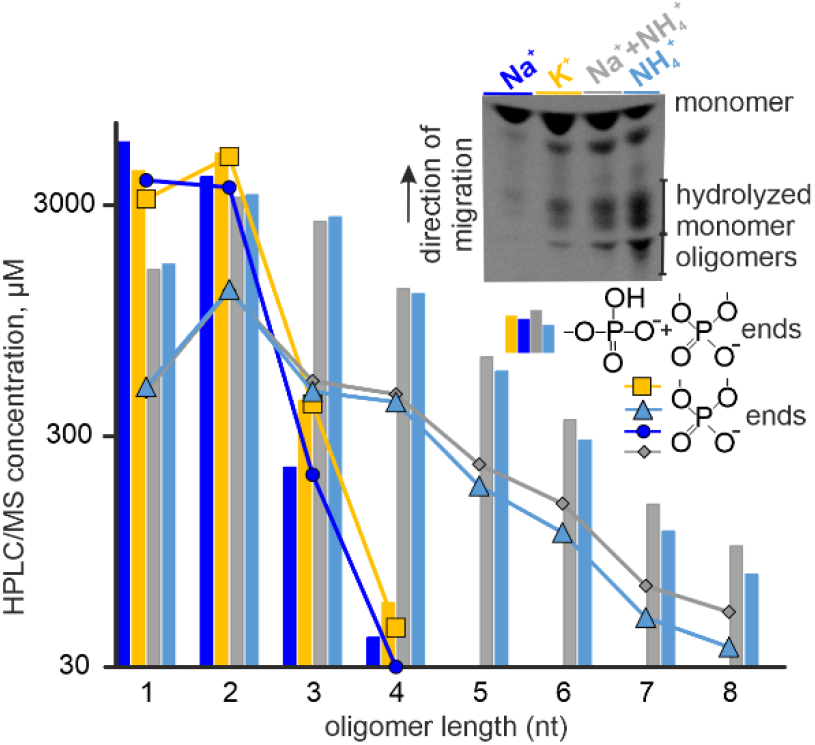
Counterion dependence. Polymerization of Na^+^, K^+^ and NH_4_^+^ forms of cCMP as well as that of a 1:1 mixture of Na^+^ and NH_4_^+^ forms of cCMP incubated at 35 °C for 2 days in a mixture of air, CO_2_ and NH_3_ generated by the thermal decomposition of ammonium carbamate at 62 °C (see the “laboratory mimic” setup in Figure 2A). The figure shows the HPLC/MS and TLC analyses (see the inset) of the reaction products. For polyacrylamide gel electrophoresis of parallel samples see Figure S3 in the Supporting Information.

To explain the observed difference between the polymerization efficiency of the studied Na^+^ and (alkyl)ammonium form of nucleotides, we performed a thermogravimetric analysis of Na^+^ and Et_3_NH^+^ forms of cCMP. The thermogravimetric curves presented in Figure 4A illustrate that for the Na^+^ form the first significant mass loss occurs predominantly at temperatures below 100 °C, i.e. in the temperature range typical to water loss. On the contrary, the Et_3_NH^+^ salt form nucleotide starts degrading at remarkably higher temperatures (mostly above 100 °C) ruling out the possibility of a simple water loss step. Loss of the highly volatile triethylamine due to thermal degradation of the Et_3_NH^+^ countercations is more consistent with these high temperatures. Thus, the complete absence of the water-loss step suggests that the dry Et_3_NH^+^ form of the cCMP material used for the polymerization experiments holds a very low water content.

**Figure 4.**
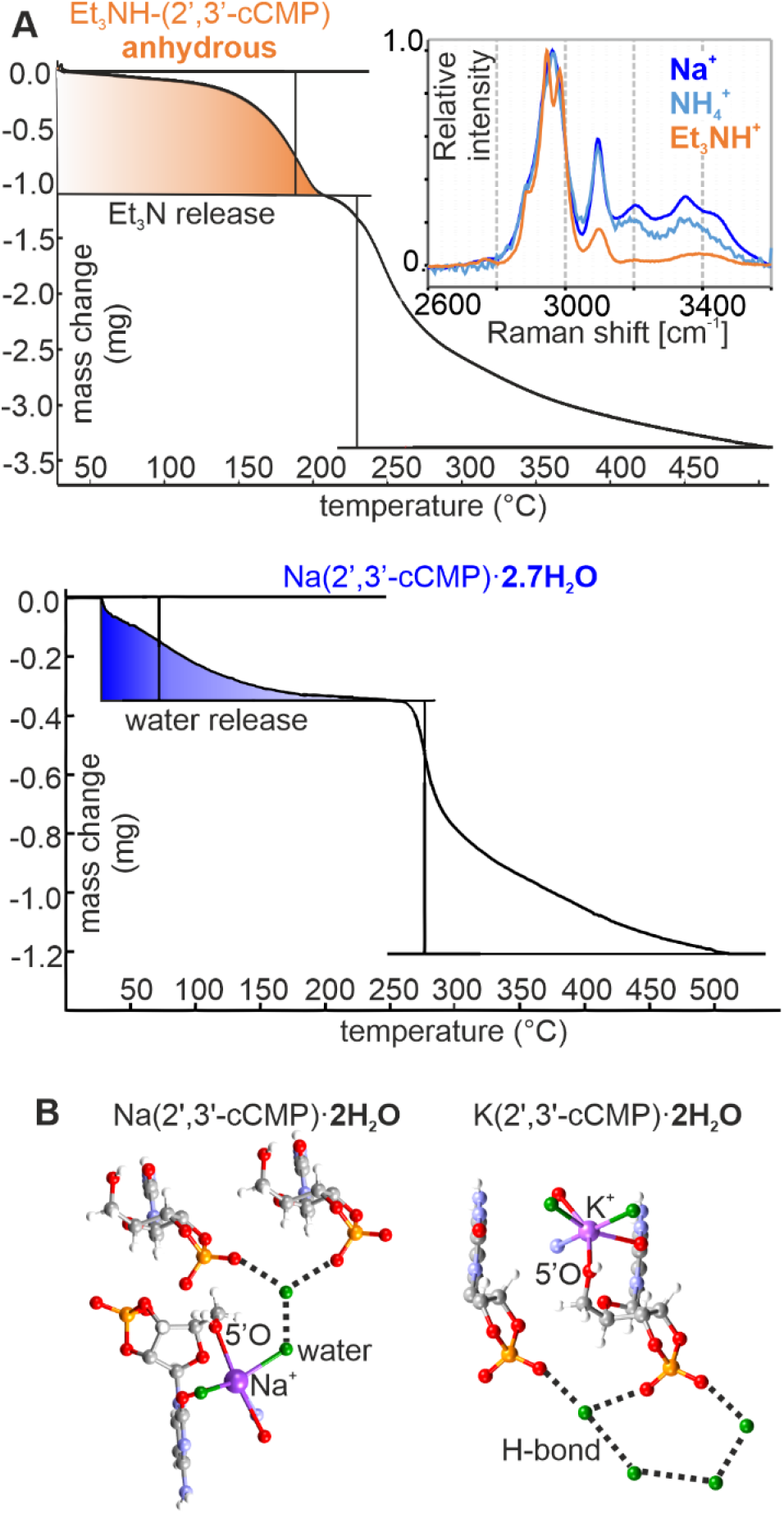
Water content of the dry material. Water content of metal salts of 2’,3’-cyclic nucleotides limits oligomer formation. A) Thermogravimetry and Raman spectra (see inset) of the amorphous Et_3_NH^+^ form of cCMP used in the polymerization experiments are consistent with a negligible water content. In contrast, based on thermogravimetry, the dry Na^+^ form material contains ca. 2.7 water molecules per nucleotide. B) Water molecules are localized in the vicinity of the 2’,3’-cyclic phosphate groups in the crystal structures of dihydrated Na^+^ (Ref. 14) and K^+^-forms of cCMP (this work) through H-bonding interactions with the anionic phosphate oxygens. In contrast, (alkyl)ammonium salts of nucleotides generally adopt an anhydrous crystalline form. Atom colors used in the nucleotide models: C−dark grey, N-blue, O−red, H−light grey, P−orange.

The above thermogravimetric analysis is further supported by our Raman spectroscopic measurements. The inset in Figure 4A compares the O-H and N-H stretching regions (3100-3600 cm^-1^)^16^ of the Raman spectra of dry Na^+^, NH_4_^+^ and Et_3_NH^+^ forms of the cCMP materials used for the polymerization experiments. In nucleic acid spectra valence vibrations of water appear as a broad band with maxima at approximately 3200, 3360 and 3430 cm^-1^.^17^ In this spectral region the normalized band intensities decrease in the order Na^+^ > NH_4+_ > Et_3_NH^+^, suggesting a remarkably higher water content for the Na^+^ form material as compared to the (alkyl)ammonium forms. In particular, the complete absence of a signal at 3300-3500 cm^-1^ in the spectrum of the Et_3_NH^+^ form nucleotide is consistent with a negligible water content of the dry material. This observation supports our proposal that binding of (alkyl)ammonium cations to the phosphate keeps water away from the transphosphorylation sites, leading to higher efficiency of the nucleotide polymerization reactions.

Molecular dynamics simulations detailed in the Supporting Information (see Figure S4) also suggest that, as compared to Na^+^ cations, NH_4+_ cations have an about 25 percent higher chance to bind to the phosphate oxygens. Thus, NH_4_^+^ cations represent potent competitors to water molecules for H-bonding with the phosphate groups of cyclic nucleotide monomers. Indeed, as Figures 2B and 3 illustrate, the NH_4_+ and Et_3_NH^+^ forms of cCMP can be polymerized into products up to octamers upon incubation in dry mixtures of ammonia and CO_2_. In contrast, efficiency of the polymerization rapidly declines for the Na^+^ and K^+^ forms (shown in Figure 3), for which most of the oligomeric products were detected in the dimer-to-tetramer region.

### Polymerization of mixtures containing cCMP, cGMP, cUMP and cAMP

In the next step, we have also attempted to polymerize two-component mixtures of NH_4_^+^ forms of cAMP with cUMP, cCMP with cGMP as well as a four-component mixture containing cCMP, cAMP, cUMP and cGMP in equimolar amounts (see Figure 5).

**Figure 5.**
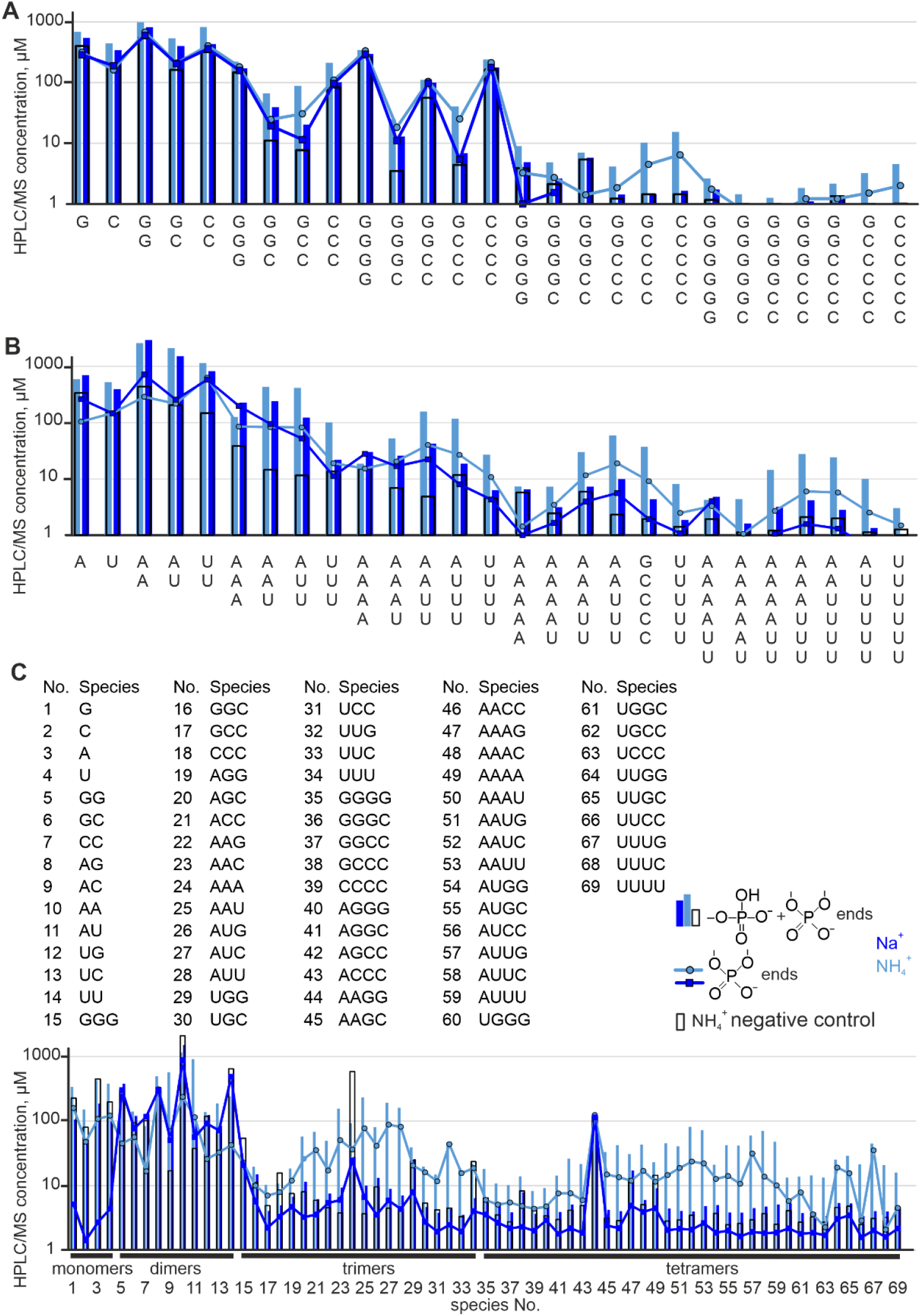
HPLC/MS analysis of copolymerization. Products obtained when polymerizing two- and four-component mixtures of cCMP, cGMP, cUMP and cAMP in NH_4_^+^ and Na^+^ forms. Sample treatment was performed at 35 °C for 2 days in a gas mixture containing air, NH_3_ and CO_2_ in the setup referred to as “laboratory mimic” in Figure 2A. Ammonium carbamate was used as a source of NH_3_ and CO_2_. A) Equimolar mixture of cCMP and cGMP. B) Equimolar mixture of cUMP and cAMP. C) Equimolar mixture of cCMP, cGMP, cUMP and cAMP. The negative control samples were prepared by incubating the corresponding mixture of NH_4_^+^ form nucleotides in the “laboratory mimic” setup by replacing the ammonium carbamate loading with water.

HPLC/MS analysis of the reaction products has shown a remarkable, 3-fold polymerization rate enhancement in the trimer-hexamer oligomer length range for the two-component mixtures containing cGMP and cCMP (Figure 5A) as well as cAMP and cUMP (see Figure 5B) in their NH_4_^+^ forms as compared to the one with Na^+^ form monomers. We note that in the mixture of cGMP and cCMP even simple drying of the nucleotides on glass without incubation in NH_3_ containing gas mixtures could trigger the polymerization, suggesting that guanine nucleotides could also serve as competent base-catalysts of the reaction. The same observation was made also for the mixture of all four nucleotides in equimolar amounts, the polymerization rate enhancement being the most remarkable (roughly 10-fold) for those sequences with higher A and U content (Figure 5C).

To extend this analysis, Figure S5 (see the Supporting Information) compares the denaturing gel electrophoretic profiles of the polymerization products formed from pure Na^+^ as well as NH_4_^+^ forms of 2’,3’-cCMP, cAMP, cUMP and cGMP. While the difference between the efficiency of the polymerization of the NH_4+_ and Na^+^ salts is relatively small for cGMP, performance of the NH_4+_ salts is clearly superior to that of the Na^+^ salts for cAMP, cUMP and cCMP. This confirms the strong rate enhancement observed for the two-component mixture containing NH_4_^+^ forms of cAMP and cUMP.

### Polymerization of the alkylammonium salts of nucleotides in a mixture of volatiles with a composition corresponding to that of the hot-water extract obtained from asteroid Bennu

To move towards a geochemically plausible polymerization scenario we have tested a mixture which contained volatiles like NH_3_, methylamine, acetic acid, formic acid and oxalic acid in a ratio found in the hot-water extract of samples taken from asteroid Bennu. Samples incubations (2 days, 35 °C) were performed in the “laboratory mimic” setup shown in Figure 2A. The HPLC/MS distribution of the polymerization products was very similar to that obtained for incubation in a vapor formed by the thermal decomposition of ammonium carbamate (Figure 6). Thus, in both cases (see the red and green bars) we see oligomer formation up to hexamers, independently on the source of the volatile amines used to initiate the polymerization reaction. This suggests that mild heating of simple organic (alkyl)ammonium salts present on extraterrestrial bodies can liberate volatile amines/NH_3_ in sufficiently high concentrations to catalyse a 2’,3’cyclic nucleotide based polymerization chemistry. For comparison, the data shown with empty bars illustrate that the polymerization reaction is clearly much less efficient for the sodium form material, despite using the same amine source (ammonium-carbamate).

**Figure 6.**
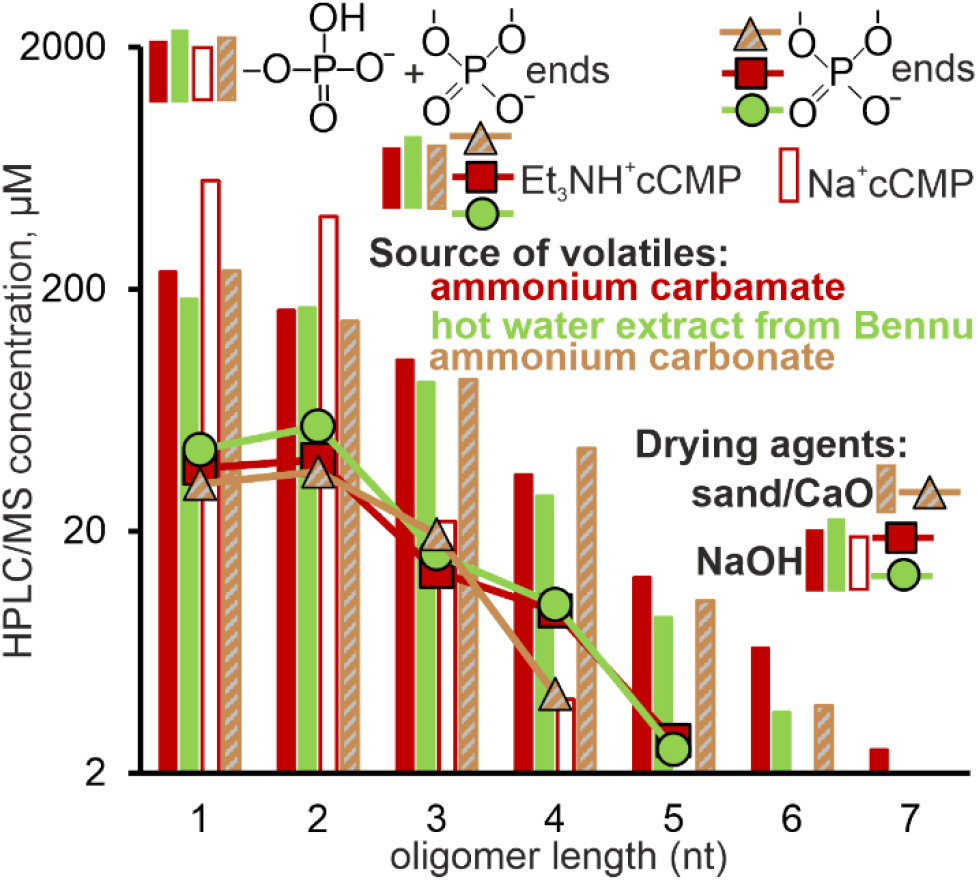
Effect of volatiles and drying agents. Length distribution of the polymerization products of Et_3_NH^+^ form of cCMP treated for 2 days in a gas mixture released from an organic mixture with a composition mimicking the hot-water extract obtained from asteroid Bennu (light green). For comparison, the data in red show the product distribution for parallel Et_3_NH^+^ and Na^+^ cCMP samples incubated in a mixture of air, NH_3_ and CO_2_ formed upon thermal decomposition of ammonium carbamate under the same conditions. In all three cases the gas mixture was dried with NaOH (see the “laboratory mimic” setup in Figure 2A). The figure also shows the product distributions observed for the polymerization of Et_3_NH^+^ form of cCMP performed in the “geological mimic” (Figure 2A) setup (brown-grey). In this latter case ammonium carbonate served as the source of NH_3_ and CO_2_, and the gas mixture was dried with a combination of a sand layer and quicklime (see the “Preparation of the samples” section in the Supporting Information).

As a final comparison, we tested the polymerization reaction in the setup referred to as “geological mimic” in Figure 2A that replaced the powerful, yet geologically less plausible, NaOH drying agent with a 5 cm thick sand layer and a thinner (ca. 1 cm) layer of CaO (quicklime) pellets on top of it. This setup was designed to ensure geologically plausible, highly efficient drying conditions that enable production of an almost water-free mixture of ammonia and CO_2_ using materials of geological origin. Porous rocks (surrogated with a sand layer) under a temperature gradient in combination with quicklime (that can easily form above 570 °C when heating Ca(OH)_2_ precipitated on the surface of volcanic rocks) seemed to be a reasonable choice for this purpose. As Figure 6 illustrates, polymerization of the Et_3_NH^+^ salt of cCMP in this setting leads to similar product composition and oligomer yields as those observed when using NaOH as drying agent.

## Discussion

### (Alkyl)ammonium charge compensating cations repel water and act as general acid-base catalysts for RNA polymerization

The TGA in Figure 4A illustrates that the water content of the dry Et_3_NH^+^ and Na^+^ forms of cCMP used for polymerization is very close to that found for the analogous crystalline salts. Whereas the crystalline NH_4_^+^ salt form is completely anhydrous,^11^ in the Na^+^ and other metal salt forms the crystal water molecules are intensively involved in direct hydrogen bonding interactions with the phosphate oxygens (Figure 4B). These water molecules are localized in the vicinity of the cyclic phosphate groups of the Na^+^ salt form cCMP monomers and are extremely well-suited to initiate hydrolytic degradation of the cyclic phosphodiester linkages of the hydrogen bond acceptor nucleotide. In contrast, ammonium salts of nucleotides often crystallize without the water of crystallization. Interestingly, crystals of the two stable (monobasic^18^ and dibasic^19^) forms of ammonium phosphate and those of the diammonium salt of hypodiphosphoric acid^20^ are completely anhydrous as well. In these crystal structures the ammonium cations serve as donors of hydrogen bonds to the phosphate oxygens. Since crystallization of ammonium phosphates from aqueous solutions involves removal of hydrogen bonded water molecules from the solvation sphere of the phosphates, this suggests that ammonium cations may efficiently compete with water molecules for binding to the phosphate oxygens.

As molecular dynamics simulations (see Figure S4 in the Supporting Information) show, ammonium charge compensating cations have a very high affinity to bind to the substrate phosphate groups through hydrogen bonding interactions. At the pK_a_ of the amines they themselves can serve as general acid-base catalysts in the polymerization reaction. The mechanism of the reaction is analogous to the one reported in our previous work^10^ for the amino acid catalyzed polymerization of 2’,3’-cyclic nucleotides (see Figure 1). Our above data demonstrate that in a volatile NH_3_/CO_2_ pH 9.2 buffer (alkyl)ammonium cations in combination with gaseous ammonia act as a very efficient catalytic pair. Thus, the (alkyl)ammonium charge compensating cations, which are excellent phosphate-binding agents, catalyze the reaction and simultaneously keep away competing water molecules from the transphosphorylation sites. These two synergic effects may explain the experimentally observed higher activity of the alkylammonium salts of 2’,3’-cyclic nucleotides as compared to their Na^+^ or K^+^ counterparts.

### Prebiotic model for the accumulation of ammonium-salt form nucleotides on the early Earth

Recent studies suggest that in pristine, extraterrestrial environments ammonium cations could serve as charge compensating species.^21-27^ For example, ammoniated phyllosilicates have been detected on the dwarf planet Ceres.^22, 27^ On our planet biotite rich in NH_4_^+^ has been found in Archean (ca. 3.8Gy old) garnet–mica schists from the Isua Supracrustal Belt in southern West Greenland.^28^ Natural ammoniated illite deposits near black shales associated with anaerobic degradation of organic matter are known from Alaska.^29^ Ammonium-rich white mica relics have been found in graphite-rich microdomains of metamorphic rocks.^30^ Ammonia has also been detected in jets of icy water plums on Enceladus.^31^

Similarly, in a prebiotic context ammonium salts could accumulate due to anaerobic degradation of organic materials, a process that is responsible for ammonia production in soil today.^32-33^ There is no question that organic tars were available in large amounts on the early Earth as a result of a non-selective organic chemistry.^34^ This material could serve as a source of ammonia and carbon dioxide. It is well-known that pyrolysis of organic tars leads to an insoluble organic matter. This process includes deamination and decarboxylation reactions that deliver large amounts of ammonia and carbon dioxide.

Pyrolysis of organic inclusions embedded in a porous rock matrix (a process still common in the Earth crust today, responsible for the formation and evolution of oil shales^35^ among others) could be induced by geothermal heat, for example due to the contact with protruding magma in volcanically active areas. This setup could also ensure temporary drying conditions that enable production of an almost water-free mixture of ammonia and CO_2_, similar to that formed in laboratory experiments using NaOH as a drying agent. Heating the porous rocks in combination with anhydrous metal-oxides derived from salt deposits (e.g. NaCl, KCl, CaCl_2_ and MgCl_2_ frequently accumulate on the surface of volcanic rocks) may provide an efficient drying medium for this purpose. A sketch of the proposed geological model and its experimental mimic is shown in Figure 2A. As our data presented in Figure 6 demonstrate, polymerization of the Et_3_NH^+^ salt of cCMP in this setting leads to similar product composition and polymerization yields as that observed using NaOH as a drying agent.

### Comparison to previous studies

We note here that early studies on the polymerization of 2’,3’-cyclic nucleotides, though without giving a reasoning, also utilized ammonium or Et_3_NH^+^ form materials, rather than Na^+^ forms.^4-5^ While Ref. 5 reported polymerization of the Et_3_NH^+^ salt of 2’,3’-cCMP up to hexamers at high-temperatures (138 °C) after a 48 hours long sample treatment, Ref. 4 demonstrated the feasibility of polymer formation from 2’,3’-cAMP at 25-85 °C upon addition of amine catalysts in at least two-fold molar excess. However, in this latter study no oligomerization was seen when excess ammonium carbonate was mixed into the reaction mixture, in contradiction to our findings. We explain this observation with the fact that in this case the sample treatment included a prolonged and prebiotically strongly questionable (oil)vacuum-drying step prior to polymerization. This step efficiently removed also the catalyst, i.e. the highly volatile ammonium-carbonate, from the sample.

## Conclusion

Besides obvious thermodynamic reasons, water interferes with prebiotic polymerization reactions of activated nucleotides mainly due to the hydrolysis of these relatively high-energy monomers.^15^ We found that by decreasing the residual water content of the dry nucleotide samples used in polymerization experiments higher polymerization yields and longer oligomers could be achieved. We used (alkyl)ammonium salts of 2’,3’-cyclic nucleotides for which, contrary to metal salts, available crystallographic data indicated that they are less likely to incorporate water in their crystal structures.^11-13^ In addition, the alkylammonium charge compensating cations can serve as general acid-base catalysts in the transphosphorylation reaction leading to inter-nucleotide phosphodiester bond formation according to an analogous mechanism as the one reported in our previous work (see Figure 1).^10^

Being a ubiquitous component of extraterrestrial materials,^21-27^ there is increasing evidence that ammonium salts could be present on the early Earth in considerable amounts. Departing from a general acid-base catalytic scenario of nucleotide polymerization, we tested the polymerization efficiency of (alkyl)ammonium salts of 2’,3’-cyclic nucleotides in combination with an ammonia-containing pH 9.2 volatile buffer. Our data demonstrate that this catalytic system benefits not only from being self-catalyzed by the charge compensating cations themselves, but also from the fact that these salts have an unusually low water content as compared to the metal salts of nucleotides. In addition, molecular dynamics simulations revealed that ammonium cations outcompete water molecules when binding to the phosphate group of cyclic nucleotides. This way they lower the chance of monomer hydrolysis that is the main deactivation pathway interfering with the studied polymerization reaction. Our analysis shows that prebiotic environments where amines or ammonia could accumulate in higher amounts could provide favorable conditions for RNA polymerization.

## Supporting information

Supplemental information to the paper by Schmid et al.

## Acknowledgements

Financial support from the GAČR/DFG Lead Agency project No. 25-16127K/549991943 as well as from Deutsche Forschungsgemeinschaft (DFG, German Research Foundation) – Project-ID 521256690 – CRC 392 and European Research Council BubbleLife #10116688, ERC-2024-SyG is greatly acknowledged.

We acknowledge CEITEC Proteomics Core Facility of CIISB, Instruct-CZ Centre, supported by MEYS CR (LM2023042, CZ.02.01.01/00/23_015/0008175). We are indebted to our colleague, Dr. Roman Matyášek, who sadly passed away during the preparation of this manuscript.

**References 36-52 refer to works cited only in the Supporting Information.

