## Supplemental information to the paper by Schmid et al. for "Enhanced RNA Formation Under Amine-Rich Local Atmospheres from 2’,3’-Cyclic Nucleotides"

by **Almuth Schmid et al.**

**Table of contents**


| | |
|---|---|
| *Experimental section* | *p. 3-6.* |
| Materials | p. 3. |
| Preparation of the samples | p. 3. |
| Thin layer chromatographic (TLC) analysis | p. 4. |
| Polyacrylamide gel electrophoresis | p. 4. |
| HPLC/MS | p. 5. |
| MALDI-ToF mass spectrometry | p. 5. |
| Thermogravimetric analysis | p. 5. |
| Raman spectroscopy | p. 6. |
| Single-crystal X-ray diffraction | p. 6. |
| *Computational methods* | *p. 7-8.* |
| System building for molecular dynamics simulations | p. 7. |
| Molecular dynamics simulations | p. 8. |
| *Supplementary figures mentioned in the manuscript* | *p. 9-13.* |
| Figure S1 | p. 9. |
| Figure S2 | p. 10. |
| Figure S3 | p. 11. |
| Figure S4 | p. 12. |
| Figure S5 | p. 13. |
| *Supplementary figures mentioned in the Supporting Information* | *p. 14-17.* |
| Figure S6 | p. 14. |
| Figure S7 | p. 15. |
| Figure S8 | p. 16. |
| Figure S9 | p. 16. |
| Figure S10 | p. 17. |
| Figure S11 | p. 17. |
| *Supplementary tables* | *p. 18-20.* |
| Table S1 | p. 18. |
| Table S2 | p. 19. |
| Table S3 | p. 20. |
| *References to the Supporting Information* | *p. 21-23.* |

### Experimental section

**Materials.** $Na^+$ salts of cytidine 2',3'-cyclic monophosphate (cCMP), guanosine 2',3'-cyclic monophosphate (cGMP), adenosine 2',3'-cyclic monophosphate (cAMP), uridine 2',3'-cyclic monophosphate (cUMP) were purchased from Biolog in lyophilized form (Cat. Nos. C104, G025, A307, U001). $NH_4^+$ salts of the same nucleotides were purchased from Biolog as custom-made 10 mM aqueous solutions.

The triethylammonium ($Et_3NH^+$) form of cCMP was synthesized according to the protocol of Veena et al.[S1] and was used as a 10 mM aqueous solution in the experiments. $^1$H-,and $^{13}$C-NMR as well as MALDI-ToF mass spectra of the synthesized material are shown in Figures S6-S8. XRD has identified leftovers of crystalline ($Et_3NH$)Cl and a small amount of crystalline nucleotide acid[S2] in the $Et_3NH^+$ form of cCMP, which cannot be avoided due to the high-vacuum treatment needed in the final purification step after ion-exchange.

$CaCl_2$ was purchased from Sigma as a crystalline dihydrate form. $(NH_3)_{aq}$ was purchased as a concentrated (22-24 wt%) aqueous solution from Penta Chemicals (Index No. 007-001-01-2). NaOH (pearls, Index No. 011-002-00-6), ammonium carbonate (Cat. No. 207861) and ammonium carbamate (Cat. No. 018134.18) were purchased from Lach-Ner, Sigma-Aldrich and Thermo Scientific, respectively. Ultrapure (Type 1) water prepared with a Direct-Q 3UV (Merck-Millipore) apparatus was used throughout the experiments.

Methylamine aqueous solution (Sigma-Aldrich, Cat. No. 426466), concentrated acetic acid (Sigma-Aldrich, Cat. No. 695092), formic acid (Sigma Aldrich, Cat. No. F0507) and oxalic acid as dihydrate (Lach-Ner Index No. 607-006-00-8) were used to prepare a suspension corresponding to the composition reported for the hot-water extract isolated from a sample taken from asteroid Bennu in Ref. S3. Composition of the mixture is given in Table S1.

CaO (quicklime) pellets were prepared by dehydration of $Ca(OH)_2$ at 650 °C for 5 hours. $Ca(OH)_2$ was prepared by precipitation from a concentrated $CaCl_2$ solution with NaOH.

**Preparation of the samples.** 10 µL of a 10 mM nucleotide solution (in total 100 nmol) was dried on glass in air flow at room temperature for 1 hour. This sample was then incubated at 35 ˚C in a tightly closed 15 mL Falcon tube in a mixture of air, $NH_3$ and $CO_2$ for 2-3 days. The length of the incubation time depended on the setup used to generate the gas mixture. In case of multicomponent equimolar mixtures of nucleotides the total amount of nucleotides used for one experiment was 100 nmol.

Two different setups were used to generate a dry air, $NH_3$ and $CO_2$ mixture. In the setup referred to as "laboratory mimic" (see Figure 2A, main text) we used a very efficient drying agent, NaOH pellets, to dry a $CO_2$ and $NH_3$-containing gas mixture that forms upon thermal decomposition of ammonium carbamate at 62 ˚C (note that ammonium carbamate always contains some carbonate as an impurity). The best performance was achieved when loading the conical part of a 15mL Falcon tube with a fine powder of ammonium carbamate (ca. 1g) covered with a 2-3 cm thick layer of NaOH pellets (separated with a teflon sheet).

In some of the experiments as a source of $NH_3$ and volatile amines we used a mixture of simple organic compounds with a composition found in the hot-water extract of a sample taken from asteroid Bennu (for compositional data see Table S1).[S3] In this case the ammonium carbonate loading was replaced with an Eppendorf tube, loaded with 750 mL of the above mixture of volatiles. The Eppendorf tube was placed into the Falcon tube and kept open in order to ensure the evaporation of its volatile content upon heating at 62 ˚C.

In the setup referred to as "geological mimic" (see Figure 2A in the main text) the NaOH drying agent was replaced with a 5 cm thick layer of commercial sand (previously burnt at 850 ˚C for 6 hours to remove its organic content) covered with a 1 cm thick layer of CaO burnt at 650 ˚C for 5 hours. The sand represented the porous rock in which a large part of the water content of the evaporates condensed, since only the bottom part of the Falcon tube (containing the source of $NH_3$) was heated. The CaO on top of the sand served as an imitation of a fresh quicklime deposit that could transiently accumulate on the early Earth due to dehydration of $Ca(OH)_2$ at temperatures around 600 ˚C. This layer served as an efficient drying agent that removed the residual water from the gas mixture. We note that $Ca(OH)_2$ could accumulate on the early Earth in larger amounts in a pH 9 environment, since $CaCl_2$ is a frequent salt deposit on rock surfaces and at this pH it is essentially converted to the much less soluble $Ca(OH)_2$. In order to mimic a situation in which the source of $NH_3$ has a high water content, in the "geological mimic" setup ammonium carbonate rather than ammonium carbamate was used as a source of $NH_3$. Note that thermal decomposition of ammonium carbonate generates equimolar amounts of carbon dioxide and water. In contrast, in ideal case, fresh ammonium carbamate decomposes into a dry mixture of $NH_3$ and $CO_2$.

**Thin layer chromatographic (TLC) analysis.** Thin layer chromatographic analyses were performed using silica-coated aluminium plates modified with a 254nm fluorescent indicator (Supelco 1.05554.001). For cytosine, uracil and guanine nucleotides the separation was performed in a running solvent containing n-propanol:$NH_{3(aq)}$(conc.):water in a 11:4:6 ratio. For adenine-nucleotides, due to their more apolar character, we have used an n-propanol:acetone:$NH_{3(aq)}$(conc.):water 10:60:3:27 mixture. Densitometric evaluation of the TLC images was performed with a Supelco TLC Scanner using the image recorded upon illumination by UV light with a wavelength of 254 nm.

**Polyacrylamide gel electrophoresis.** Products of polymerization reactions were labelled at the 5' terminus with γ[$^{32}$P] ATP in a polynucleotide kinase (PNK) reaction. Briefly, polymerization products were precipitated by addition of sodium acetate pH 5.2 to a final concentration of 0.3 M and 2.5 volumes of ice-cold ethanol. After incubation at -20 °C for 2 hours the precipitate was sedimented by centrifugation (15 000 rpm, 5 min) and dissolved in 15 µL of distilled water. The 50 µL labelling reaction mixture contained 15 µL polymerization products, 5.0 µL 10× PNK buffer, 1.0 µl (10 µCi) γ[$^{32}$P]-ATP (Hartmann analytic, Braunschweig, Germany) and 1.0 µL (10 units) PNK (NEB, Lithuania) and 38 µL distilled water. The reaction was carried out at 37 °C for 60 min and terminated by the addition of 180 µL $H_2O$ and 200 µL phenol-chlorophorm-isoamylalcohol (25:24:1) mixture. Oligonucleotides were then extracted into the water phase by vortexing and centrifugation in an Eppendorf tube (15000 rpm, 1 min). The upper phase was removed and oligonucleotides were precipitated with sodium acetate (pH 5.2, added to a final concentration of 0.3 M), followed by the addition of 2.5 volumes of ice-cold absolute ethanol and 1 µg glycogen (R0561, Thermo Fisher Scientific, Waltham, USA). After incubation at -20 °C for 2 hours the precipitate was sedimented by centrifugation (15000 rpm, 5 min). The sediment was washed with absolute ethanol and air-dried. The radioactively labeled fragments were dissolved in the electrophoretic buffer containing 100% formamide and bromphenol blue and separated on a 15% polyacrylamide gel (Hoefer Scientific (USA) apparatus). Size markers included $C_3$ and $C_{24}$ RNA oligomers (Biomers, Germany) radioactively labelled with γ[$^{32}$P]-ATP as above. The gel running conditions were as follows: 1× TBE electrophoretic buffer, 55 °C, 35 cm glass length and run length of about 5 hours. The radioactivity of the gels was scanned using a Phosphorimager (Typhoon 7300, GE Healthcare, England) and bands were visualized by the ImageQuant (GE Healthcare) software. Exposure times to phosphorscreens were typically 15-30 min.

**HPLC/MS.** Oligomer quantification was performed using high-performance liquid chromatography (Agilent 1260 Infinity II) coupled with an electrospray ionisation time-of-flight mass spectrometer (Agilent 6230B with Dual AJS ESI). To ensure accuracy and determine the timing of the HPLC, pre-formed oligomers (polyC) ranging from lengths 2-10 with 3' phosphate (from Biomers) were used as standards. Reverse-phase ion-pairing HPLC was employed to separate oligomers of different lengths on an Agilent AdvanceBio Oligonucleotide C18 Column (4.6 x 150 mm, 2.7 µm, heated to 60 °C), with a gradient elution at a flow rate of 1 mL/min. The eluents consisted of water (Bottle A) and 50:50 methanol-water mixture (Bottle B) containing 8 mM triethylamine (TEA) and 200 mM hexafluoroisopropanol (HFIP). The separation process began with a 5-minute flush with 1% B, followed by a gradual increase to 30% B over 22.5 minutes and then to 40% B over 15 minutes. Subsequently, the column was flushed with 100% B for 5 minutes before re-equilibration at 1% B for 6 minutes. Detection of eluted oligonucleotides was achieved using ESI-TOF in negative mode, employing specific source parameters: Gas temperature: 325 °C, Drying gas flow: 13 L/min, Sheath gas temperature: 400 °C, Sheath gas flow: 8 L/min, VCap: 3500V, Nozzle Voltage: 2000V.

**MALDI-ToF mass spectrometry.** The samples were mixed with the MALDI matrix (3-hydroxypicolinic acid, 75 mg/mL in 20 mg/mL dibasic ammonium citrate:acetonitrile, 1:1 v/v mixture) in 1:4 v/v ratio. After being applied to a stainless steel sample target, the samples were analyzed in reflectron positive ion detection mode. The analyses were performed with an Ultraflextreme MALDI-ToF mass spectrometer (Bruker Daltonics, Bremen, Germany).

**Thermogravimetric analysis.** Thermogravimetric TG/DTA analysis for 2',3'-cCMP sodium salt (m = 2.748 mg) was performed on a SETARAM SETSYS 16/18 instrument in the temperature range of 30-500 °C at a heating rate of 10 °C/min, under a nitrogen flow. TG/DSC measurement for 2',3'-cCMP triethylammonium salt (m = 5.994 mg) was performed with the use of a Mettler-Toledo TGA/DSC 3+ instrument in the temperature range 30-500 °C with a ramp rate of 10 °C/min, in flowing nitrogen (flow rate: 3 $dm^3/h$).

**Raman spectroscopy.** Raman spectra of dry $Na^+$, $NH_4^+$ and $Et_3NH^+$ form cCMP samples were obtained using an upright confocal Raman microscope WITec alpha300 RSA (Oxford Instruments - WITec, Germany) equipped with the 50× dry objective EC Epiplan-Neofluar LD DIC, NA 0.55 (Zeiss, Germany) and optical-fiber-connected spectrograph UHTS300 (Oxford Instruments - WITec, Germany) with a Newton 970 EMCCD camera (Oxford Instruments - Andor, Ireland) optimized for detection in the visible region. The 532 nm laser with the excitation power of approximately 20 mW at the focal plane was used.

All samples were analysed as thin films deposited on a glass slide. Since the film thickness was less than 5 μm, their Raman spectra were extracted from axial scans of dimensions 80 × 8 μm (length × depth), where the scanned depth section covered the entire film thickness as well as a part of the glass slide. The scanning step in the sample plane was 500 nm, in the axial direction 200 nm, with an integration time of 100 ms per voxel. This method of measuring Raman spectra enabled the monitoring of the sample homogeneity and distinguishing the spectral contribution from the glass slide. Raman scans were acquired from 5 to 10 different locations on each sample and analyzed using the WITec Project Six Plus ver. 6.2 software (Oxford Instruments - WITec, Germany) by implementing advanced cosmic ray removal, background subtraction, and spectral demixing with the True Component Analysis tool (Oxford Instruments - WITec, Germany). Using the True Component Analysis tool, a representative spectrum was generated for each Raman scan, characterizing only the cytosine layer and excluding the glass-slide contribution. Spectra obtained from different locations of the same sample were compared, and the resulting spectrum was generated as the average of these spectra.

**Single-crystal X-ray diffraction.** X-ray quality crystals of K(2',3'-cCMP)·$2H_2O$ were prepared by seeding a concentrated solution of the dissolved monopotassium salt with Na(2',3'-cCMP)·$2H_2O$ (obtained by slow diffusion of ethanol into the aqueous solution of the salt). The crystals of K(2',3'-cCMP)·$2H_2O$ were then grown by slow diffusion of ethanol vapor into the saturated solution. Diffraction data for the crystal were collected on a Rigaku κ-geometry XtaLAB Synergy DW four-circle diffractometer (rotating anode X-ray source, Cu Kα radiation, ω scan method) with a hybrid HyPix-Arc 150° detector, at 100 K. Data were corrected for Lorentz and polarization effects and absorption (by the empirical (multi-scan) method; see Table S2 for details). Data collection, processing, and analysis were carried out with CrysAlis PRO.[S4] The structure was solved using dual-space algorithm with the SHELXT program.[S5] and refined on $F^2$ by a full-matrix least-squares technique using the SHELXL program,[S6] with anisotropic displacement parameters for non-H atoms.

Hydrogen atoms were located in difference Fourier maps. The H atoms from cytosine $NH_2$ groups and ribose O5'(H) groups were first refined freely with isotropic displacement parameters. Then the restraints on the N−H and O–H bonds (0.880(2) Å) and 0.840(2) Å, respectively) were applied. In the final refinement steps the H atoms coordinates were constrained to ride on the parent (N or O) atoms (AFIX 3 instructions in SHELXL). The H atoms from CH and $CH_2$ groups were refined using a riding model, with C–H = 0.95-1.00 Å, and with $U_{iso}(H) = 1.2U_{eg}(C)$. Hydrogen atoms of water molecules were isotropically refined at first, then the restraints on the O−H bonds (0.840(2) Å) and H−H distances (1.360(2) Å) were applied, and finally the rigid body constraints were used (AFIX 6).

The potassium salt is isostructural with the sodium analogue, Na(2',3'-cCMP)·$2H_2O$,[S7] therefore for K(2',3'-cCMP)·$2H_2O$ a similar asymmetric unit was chosen (Figure S9). The coordinates were then moved by a vector (0.5, 0, 0.5).

Details of structure refinement are given in Table S2. Structural data, like characteristic hydrogen-bonding contacts are listed in Table S3. Coordination environment of the $K^+$ cations and crystal packing are shown in Figures S10 and S11. The crystallographic information file (CIF) is deposited at the Cambridge Crystallographic Data Centre (CCDC No. 2523543 ) and provided as Supporting Information. The DIAMOND program was used to make the figures.[S8]

### Computational methods

**System building for molecular dynamics simulations.** The parameters for the cytidine 2',3'-cyclic monophosphate (cCMP) were taken from an earlier work[S9] and its behavior was described using the OL3 RNA force field.[S10] The ammonium cation ($NH_4^+$) parameters in AMBER do not contain the explicit hydrogens. Therefore, to obtain parameters of the complete $NH_4^+$ cation for our simulations, we first calculated the electrostatic potential of the $NH_4^+$ cation using the Gaussian 09 (rev. A02) and HF/6-31G* level of theory. The partial charges were subsequently fitted using the RESP procedure.[S11] The final partial charges obtained were -0.708421 on the nitrogen atom and 0.427105 on each of the four hydrogens. The vdW radii, bond lengths and angles for the $NH_4^+$ cation were taken from the ff14SB AMBER force field for proteins.[S12] To prevent artificial clustering of the cCMP molecules in simulations, we additionally applied the stafix modification with a factor of 0.5.[S13]

Two types of simulation systems were subsequently constructed using the xLeap module of AMBER 22:[S14] a) systems containing five cCMPs along with either five $Na^+$ or five $NH_4^+$ cations, and b) a system containing four cCMPs along with a mixture of two $Na^+$ and two $NH_4^+$ cations. The initial spatial arrangement of the cCMP molecules and counterions was randomly selected during system preparation (Figure S4), ensuring no steric overlaps. Each system was then solvated in an octahedral box of SPC/E water molecules, with minimum solute-to-box-edge distances of 8 Å and 7 Å for the systems containing five and four cCMP molecules, respectively. These box dimensions were determined through trial-and-error as the smallest configurations that avoided finite-size artifacts in periodic boundary conditions while maintaining computational efficiency.

**Molecular dynamics simulations.** Energy minimization and equilibration were carried out using pmemd.MPI in AMBER 22, following previously established procedures.[S15] Production simulations were then performed for 5 μs for each of the systems, using pmemd.cuda on RTX 3080 Ti GPUs. Two independent simulations of each system were performed using separate equilibration stages and unique random seeds assigned to the initial velocity distributions. All simulations employed SHAKE[S16] constraints in combination with hydrogen mass repartitioning,[S17] allowing for a 4 fs integration timestep. Long-range electrostatics were treated using the particle mesh Ewald (PME)[S18] method under periodic boundary conditions, with an 8 Å cutoff for Lennard-Jones interactions. System temperature and pressure were controlled using a Langevin thermostat and a Monte Carlo barostat, respectively. The production simulations were subsequently visualized in VMD[S19] and their analysis performed in cpptraj.[S20] We employed the hbond command in cpptraj to evaluate the occupancies of $Na^+$ and $NH_4^+$ cations in the vicinity of the phosphate group of cCMP. Cations were considered bound when the distance between their heavy atom and any atom of the phosphate group was less than 3.5 Å. Total occupancies of $Na^+$ and $NH_4^+$ across all cCMP molecules in the system were summed, and the relative binding preference between the two cations was determined by taking the ratio of these summed occupancies.

**Supplementary figures mentioned in the manuscript**

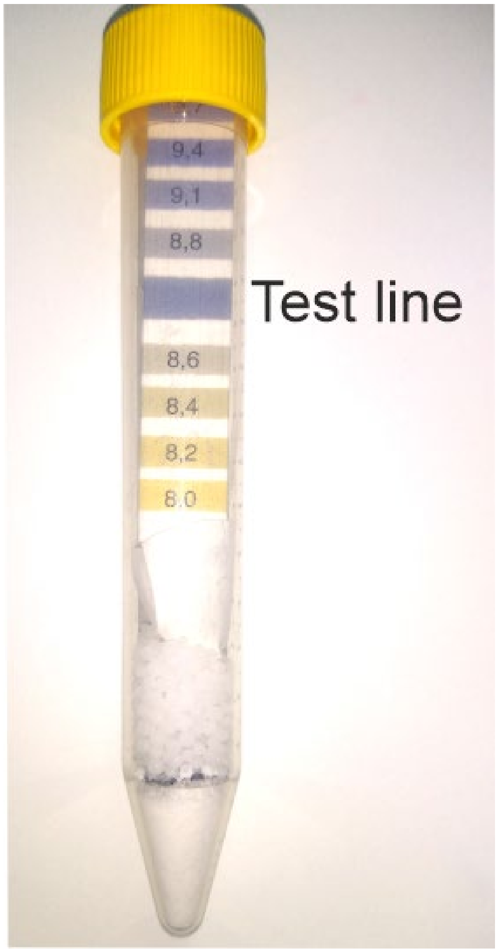


**Figure S1. pH of the gas-phase above ammonium-carbamate heat-treated at 62 °C is 9.2.** "Laboratory mimic" setup (shown in Figure 2A, main text).

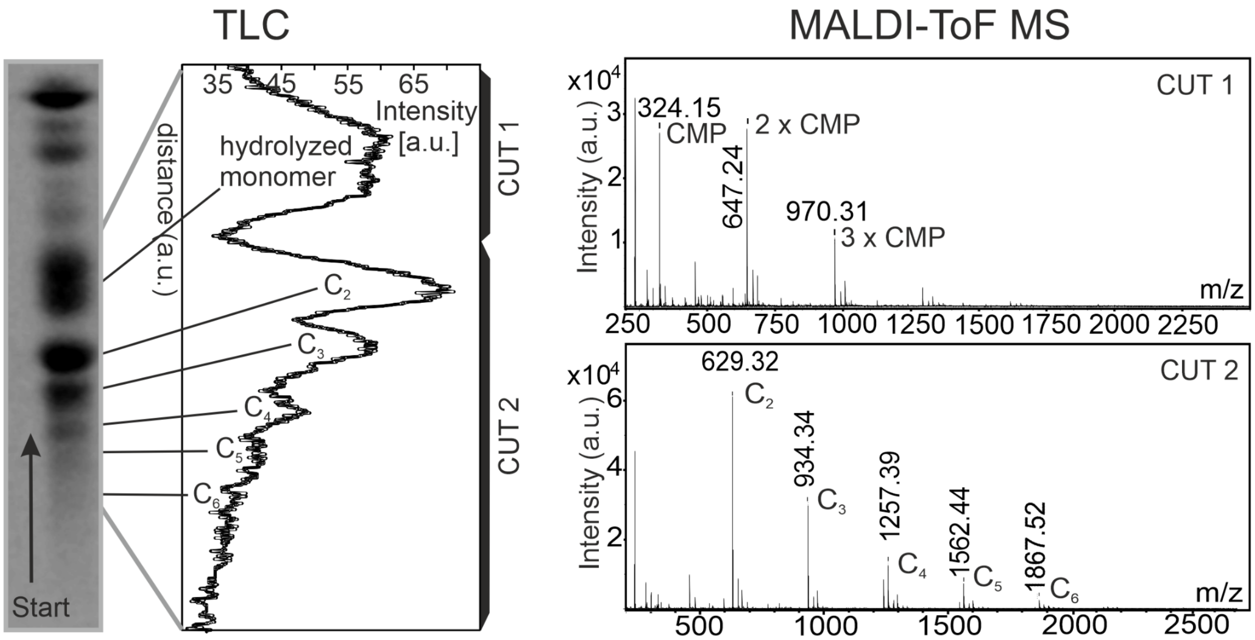


**Figure S2. MALDI-ToF MS analysis of the materials isolated (labelled as “CUT 1” and “CUT 2”) from the thin layer chromatographic bands of the polymerized $Et_3NH^+$ salt form cCMP material shown in Figure 2, main text.** $C_n$ refers to n-mer cytidylic acid oligomers.

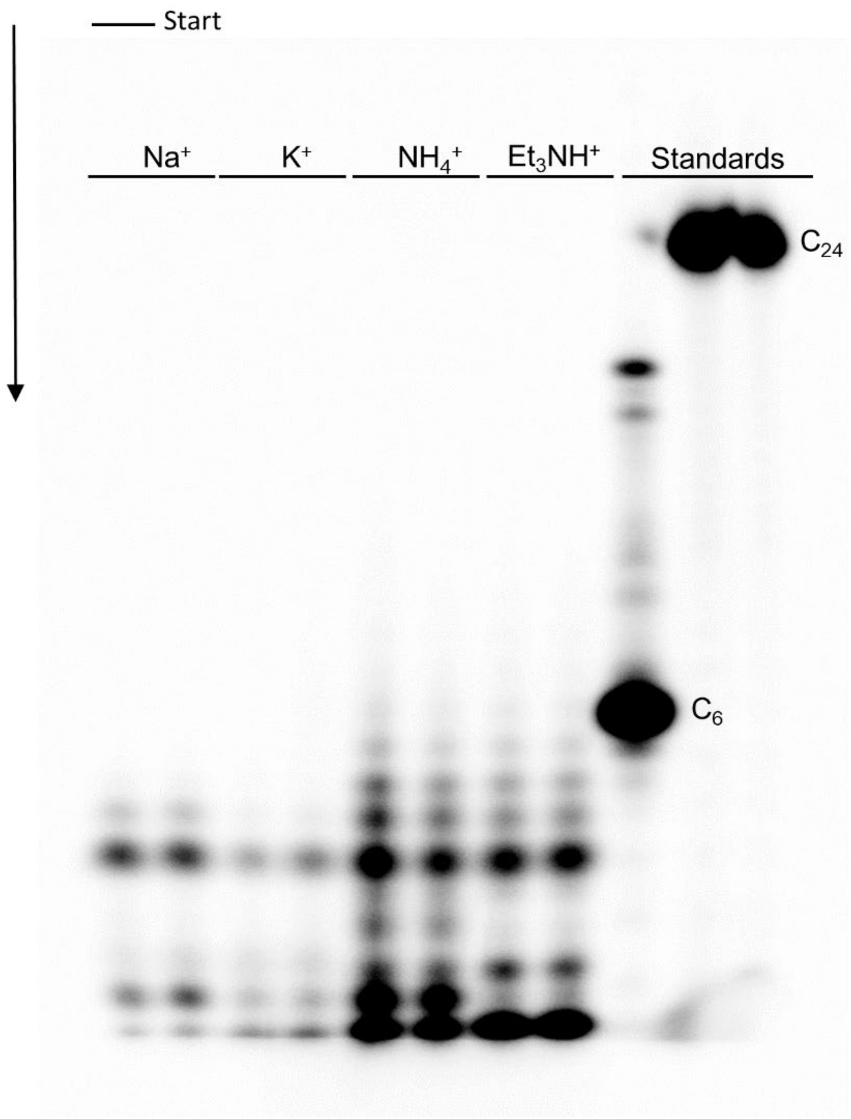


**Figure S3. Polyacrylamide gel electrophoretic analysis of oligomeric products formed from $Na^{+}$, $K^{+}$, $NH_4^{+}$ as well as from $Et_3NH^{+}$ salt forms of cCMP.** Conditions of sample preparation were the same as for the samples presented in Figure 3, main text.

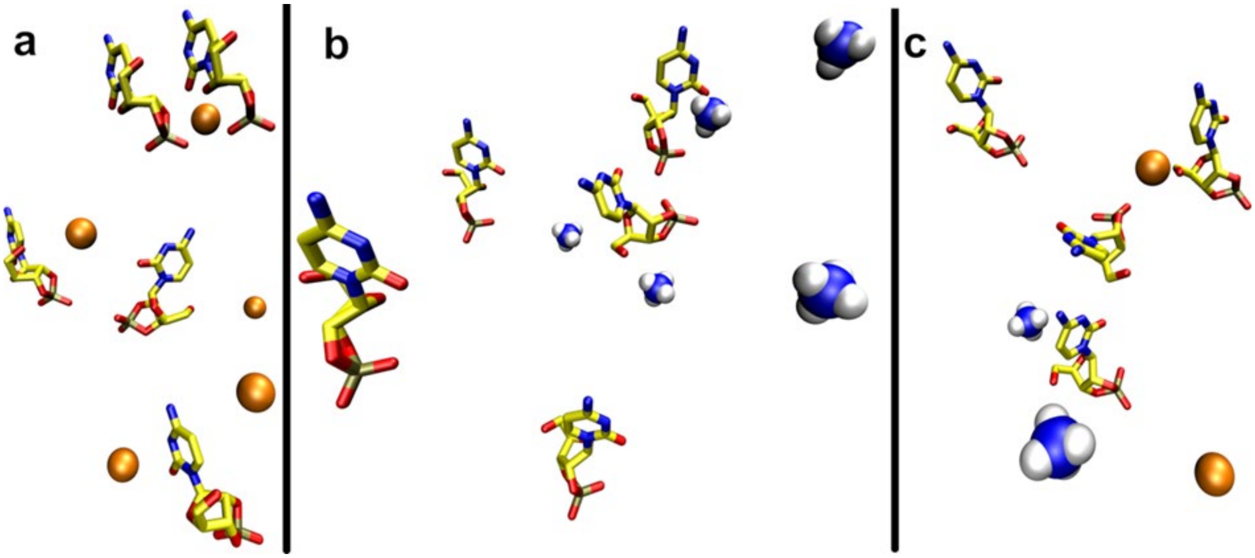


**Figure S4. Starting structures of the MD simulation systems: (a) five cCMP molecules with $Na^+$ counterions, (b) five cCMP molecules with $NH_4^+$ counterions, and (c) four cCMP molecules with an equimolar mixture of $Na^+$ and $NH_4^+$ counterions.** Hydrogen, carbon, oxygen, phosphorus, and sodium atoms are shown in white, yellow, red, bronze, and orange, respectively. For clarity, nucleotide hydrogen atoms are omitted. Based on the simulation data we have found that in the systems containing five $NH_4^+$ ions, the counterions interacted with the phosphate groups on average 24% more frequently than in systems containing five $Na^+$ ions. This difference became even more pronounced in systems containing an equimolar mixture of $Na^+$ and $NH_4^+$, where the $NH_4^+$ exhibited a 33% higher affinity for the phosphate groups compared to $Na^+$. Since $NH_4^+$ competes with water for binding to the phosphate through hydrogen bonding interactions, it essentially helps at removing water from the vicinity of the phosphate groups. In spite of the 100-fold excess of water molecules relative to the number of countercations in the simulated system, the higher affinity of the phosphate group to bind the $NH_4^+$ cations has noticeably influenced the frequency of water-phosphate binding contacts. As compared to the system with $Na^+$ countercations, the frequency of water binding to the phosphate was reduced to 98.3 and 96.7% in the 1:1 $NH_4^+$:$Na^+$ mixture and in the pure $NH_4^+$ containing systems, respectively.

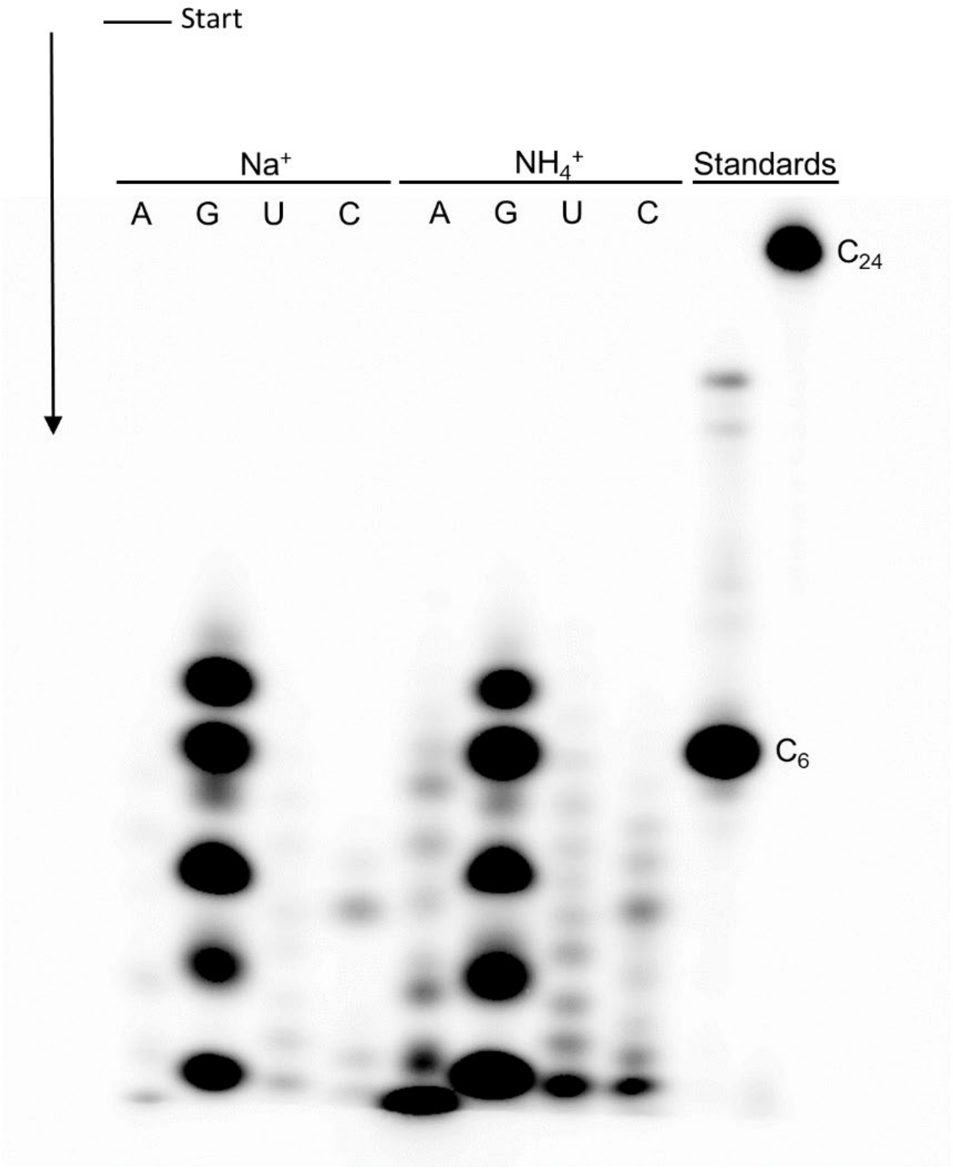


**Figure S5. Polyacrylamide gel electrophoretic analysis of $Na^+$ and $NH_4^+$ salt forms of cCMP, cUMP, cAMP and cGMP samples treated at 35 °C for 2 days in the setting referred to as "laboratory mimic" in Figure 2A, main text.** The $NH_3$- and $CO_2$- containing gas mixture was generated by the thermal decomposition of ammonium carbamate.

**Supplementary figures mentioned in the Supporting Information**

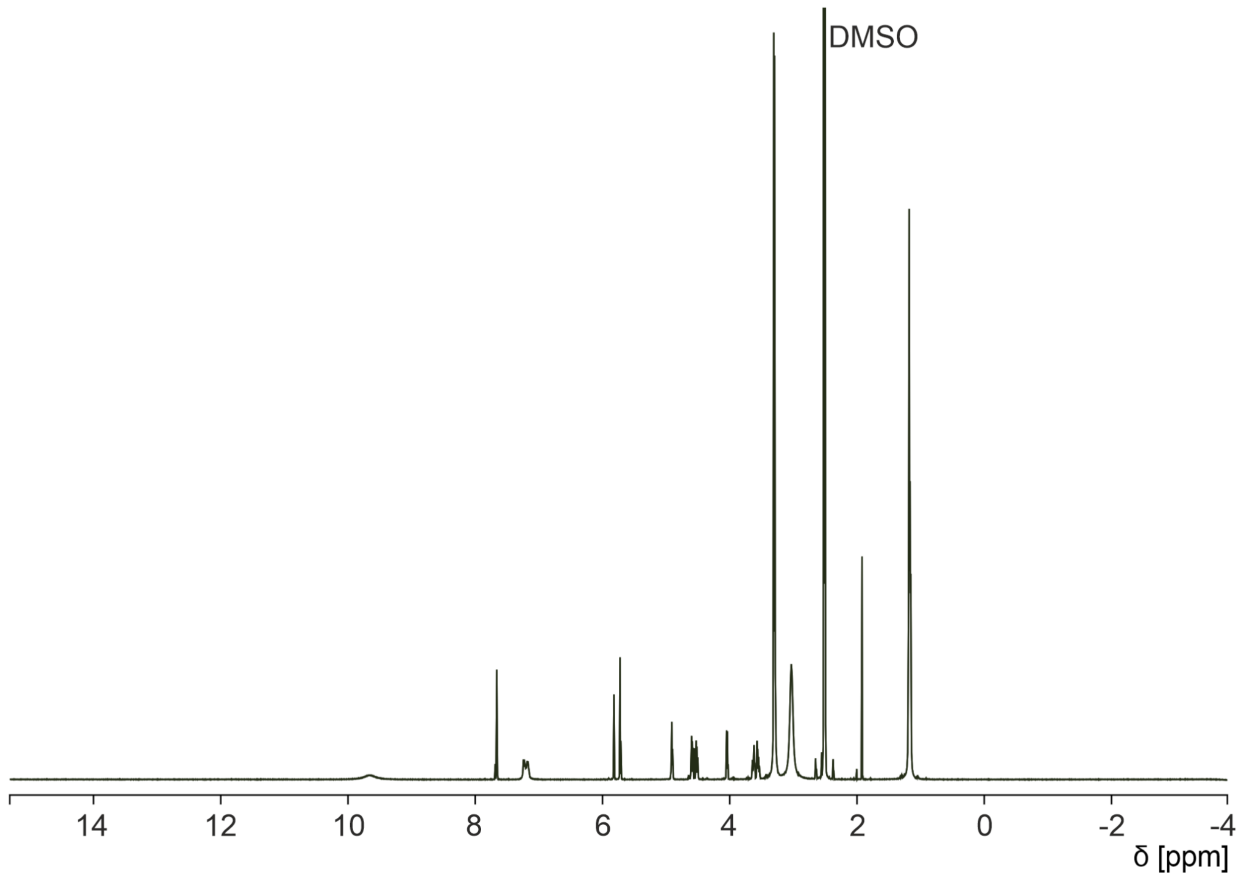


**Figure S6. $^1$H NMR spectrum (DMSO) of the synthesized $Et_3NH^+$ salt form of cCMP used in the polymerization experiments.**

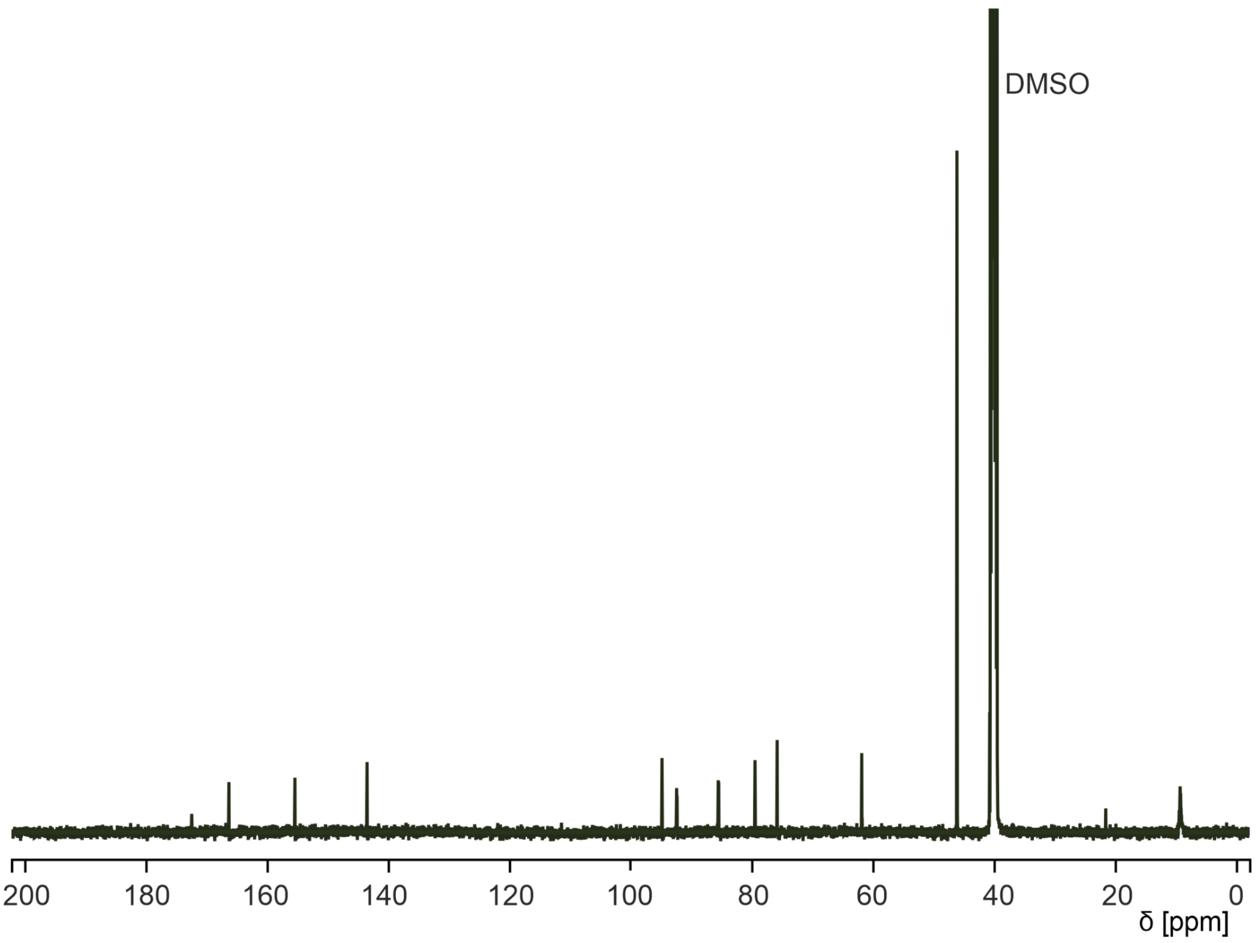


**Figure S7. $^{13}C$ NMR spectrum (DMSO) of the synthesized $Et_3NH^+$ salt form of cCMP used in the polymerization experiments.**

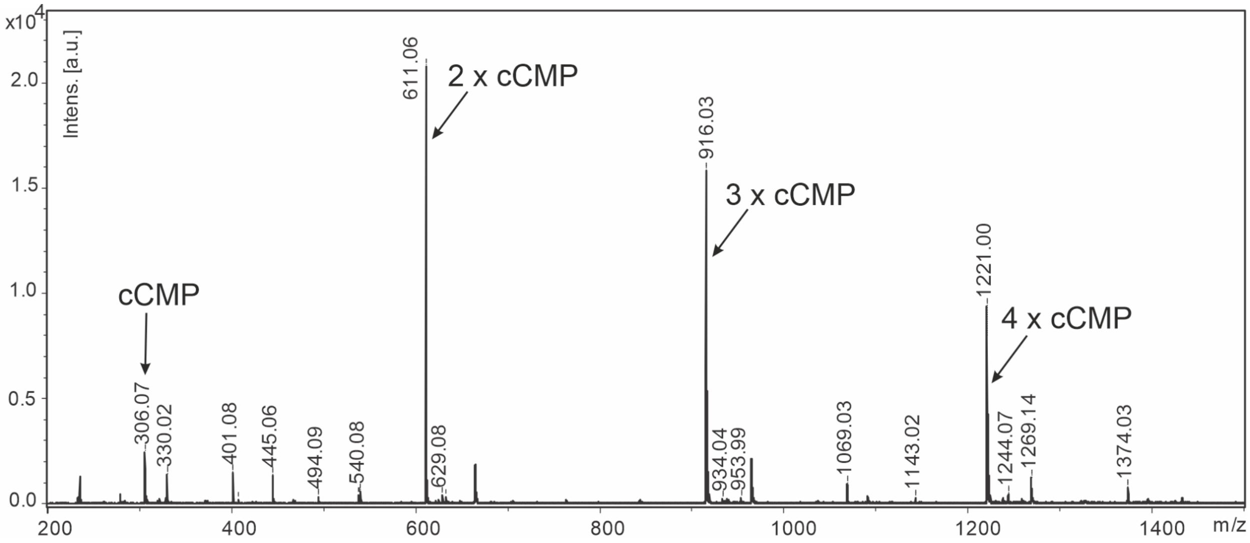


**Figure S8. MALDI-ToF mass spectrum (positive ion detection mode) of the synthesized $Et_3NH^+$ salt form of cCMP used in the polymerization experiments.**

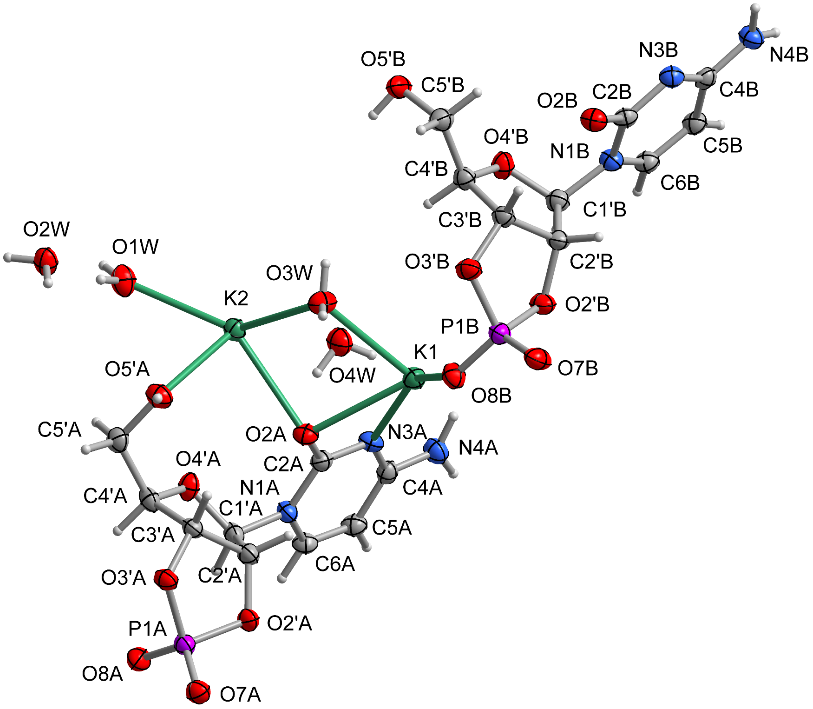


**Figure S9. Asymmetric unit of the K(2',3'-cCMP)·2$H_2O$ crystal showing the atom-numbering scheme.** Displacement ellipsoids are shown at 50% probability level.

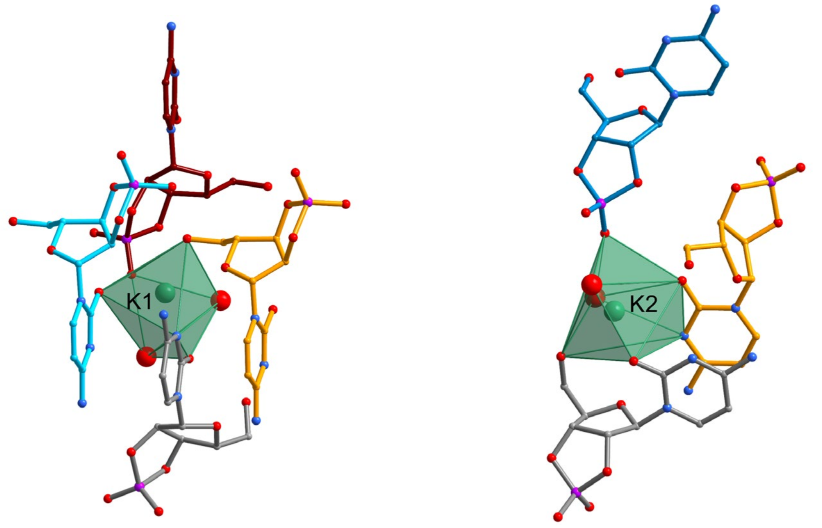


**Figure S10. Coordination environment of the $K^+$ cations (shown as green polyhedra; CN = 7) in the K(2',3'-cCMP)·$2H_2O$ crystal.** Large red balls represent water molecules.

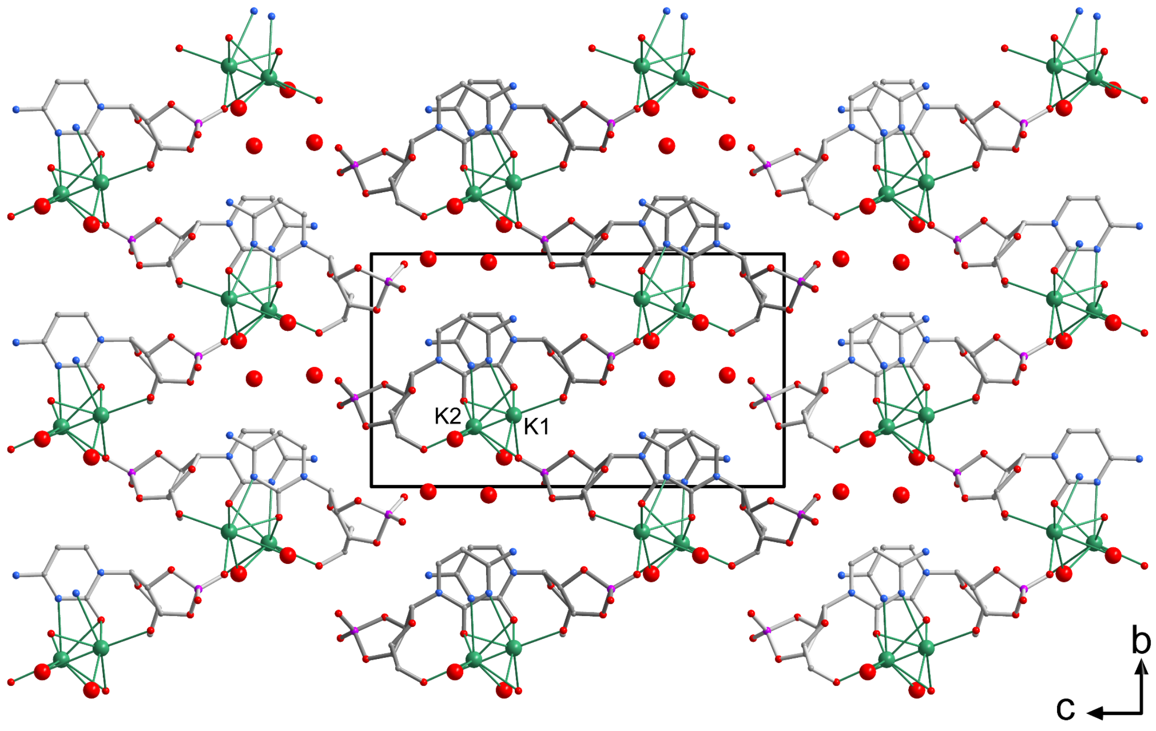


**Figure S11. Crystal packing of the K(2',3'-cCMP)·$2H_2O$ crystal.** Colour scheme: K, green; P, purple; O, red; N, blue; C, grey. H atoms not shown. Large red balls represent water molecules.

**Supplementary Tables**

**Table S1. Composition of the mixture used as a mimic of the hot-water extract isolated from a sample taken from asteroid Bennu (based on Ref. S3)**

| **Component** | **Amount** |
|---|---|
| $NH_4OH$* | 136 mmol |
| methylamine | 9.1 mmol |
| formic acid | 41 mmol |
| acetic acid | 14 mmol |
| oxalic acid | 8 mmol |

*Added as $(NH_3)_{aq}$(conc.).

**Table S2. Crystal data for K(2',3'-cCMP)·$2H_2O$.**

| | |
|---|---|
| Chemical formula | $K(C_9H_{11}N_3O_7P)\cdot 2H_2O$ |
| CCDC No. | 2523543 |
| $M_r$ | 379.31 |
| Crystal system, space group | Monoclinic, $P2_1$ |
| Temperature (K) | 100 |
| *a*, *b*, *c* (Å) | 6.8971(9), 11.0602(15), 19.656(3) |
| *β* (°) | 93.15(2) |
| *V* ($Å^3$) | 1497.2(4) |
| *Z* | 4 |
| Radiation type | Cu *K*α |
| μ ($mm^{-1}$) | 4.64 |
| Crystal size (mm) | 0.08 × 0.05 × 0.04 |
| Diffractometer | Rigaku XtaLAB Synergy DW system, HyPix-Arc 150° |
| Absorption correction | Multi-scan |
| $T_{min}$, $T_{max}$ | 0.690, 1.000 |
| No. of measured, independent and observed [$I > 2\sigma(I)$] reflections | 42314, 5967, 5700 |
| $R_{int}$ | 0.042 |
| $(\sin \theta/\lambda)_{max}$ ($Å^{-1}$) | 0.629 |
| $R[F^2 > 2\sigma(F^2)]$, $wR(F^2)$, $S$ | 0.032, 0.084, 1.10 |
| No. of reflections, parameters, restraints | 5967, 437, 19 |
| $\Delta\rho_{max}$, $\Delta\rho_{min}$ (e $Å^{-3}$) | 0.27, −0.25 |
| Absolute structure parameter | −0.002(4) |

Computer programs: *CrysAlis PRO* (Rigaku OD, 2024), *SHELXT-2014/7* (Sheldrick, 2015),*SHELXL2016*/6 (Sheldrick, 2015).

**Table S3**. **Hydrogen-bond geometry (Å, °) for K(2',3'-cCMP)·2$H_2$O.**

| *D*—H⋯*A* | *D*—H | H⋯*A* | *D*⋯*A* | *D*—H⋯*A* |
|---|---|---|---|---|
| N4*A*—H4*AA*⋯O4*W*$^{iv}$ | 0.88 | 2.36 | 3.122 (5) | 145 |
| N4*A*—H4*AB*⋯O4′*B*$^{i}$ | 0.88 | 2.49 | 3.136 (5) | 131 |
| C5*A*—H5*A*⋯O4*W*$^{iv}$ | 0.95 | 2.35 | 3.118 (5) | 137 |
| C6*A*—H6*A*⋯O3′*A*$^{v}$ | 0.95 | 2.44 | 3.307 (5) | 152 |
| O5′*A*—H5′*A*⋯O8*A*$^{vi}$ | 0.84 | 1.87 | 2.698 (4) | 170 |
| C2′*A*—H2′*A*⋯O1*W*$^{i}$ | 1.00 | 2.44 | 3.302 (5) | 144 |
| N4*B*—H4*BA*⋯O4′*A*$^{vii}$ | 0.88 | 2.51 | 3.129 (4) | 128 |
| N4*B*—H4*BB*⋯O7*A*$^{x}$ | 0.88 | 2.01 | 2.854 (5) | 162 |
| O5′*B*—H5′*B*⋯O7*B*$^{iii}$ | 0.84 | 1.89 | 2.699 (4) | 163 |
| C4′*B*—H4′*B*⋯O7*B*$^{iii}$ | 1.00 | 2.48 | 3.244 (5) | 132 |
| C5′*B*—H5′4⋯O2*B* | 0.99 | 2.60 | 3.302 (5) | 128 |
| O1*W*—H1*W*⋯O2*W* | 0.84 | 1.99 | 2.817 (4) | 170 |
| O1*W*—H2*W*⋯O2*A*$^{iii}$ | 0.84 | 2.01 | 2.766 (4) | 149 |
| O2*W*—H3*W*⋯O7*A*$^{vi}$ | 0.84 | 1.99 | 2.827 (4) | 174 |
| O2*W*—H4*W*⋯O8*A*$^{ix}$ | 0.84 | 1.97 | 2.792 (4) | 164 |
| O3*W*—H5*W*⋯O7*B*$^{iii}$ | 0.84 | 1.91 | 2.752 (4) | 176 |
| O3*W*—H6*W*⋯O4*W* | 0.84 | 1.96 | 2.792 (5) | 170 |
| O4*W*—H7*W*⋯O2*W*$^{ii}$ | 0.84 | 2.06 | 2.893 (4) | 174 |
| O4*W*—H8*W*⋯O8*B* | 0.84 | 1.93 | 2.758 (4) | 168 |

Symmetry codes: (i) −*x*+1, *y*+1/2, −*z*+1; (ii) *x*−1, *y*, *z*; (iii) *x*+1, *y*, *z*; (iv) *x*, *y*+1, *z*; (v) −*x*+1, *y*+1/2, −*z*+2; (vi) −*x*+1, *y*−1/2, −*z*+2; (vii) −*x*+1, *y*−1/2, −*z*+1; (x) *x*, *y*, *z*−1; (ix) −*x*+2, *y*−1/2, −*z*+2.

**References to the Supporting Information:**


S1. Veena, K. S.; Cruz, H. A.; Krishnamurthy, R. Microwave-Assisted One-Step Synthesis of 2′,3′-Cyclic Phosphates of Nucleosides. *Curr. Protoc.* **2023**, *3* (7), e834. https://doi.org/10.1002/cpz1.834

S2. Reddy, B. S.; Saenger, W. Molecular and Crystal Structure of the Free Acid of Cytidine 2',3'-Cyclophosphate. *Acta Crystallogr. Sect. B* **1978**, *34* (5), 1520-1524. https://doi.org/10.1107/S0567740878006032

S3. Glavin, D. P.; Dworkin, J. P.; Alexander, C. M. O. D.; Aponte, J. C.; Baczynski, A. A.; Barnes, J. J.; Bechtel, H. A.; Berger, E. L.; Burton, A. S.; Caselli, P.; Chung, A. H.; Clemett, S. J.; Cody, G. D.; Dominguez, G.; Elsila, J. E.; Farnsworth, K. K.; Foustoukos, D. I.; Freeman, K. H.; Furukawa, Y.; Gainsforth, Z.; Graham, H. V.; Grassi, T.; Giuliano, B. M.; Hamilton, V. E.; Haenecour, P.; Heck, P. R.; Hofmann, A. E.; House, C. H.; Huang, Y.; Kaplan, H. H.; Keller, L. P.; Kim, B.; Koga, T.; Liss, M.; McLain, H. L.; Marcus, M. A.; Matney, M.; McCoy, T. J.; McIntosh, O. M.; Mojarro, A.; Naraoka, H.; Nguyen, A. N.; Nuevo, M.; Nuth, J. A.; Oba, Y.; Parker, E. T.; Peretyazhko, T. S.; Sandford, S. A.; Santos, E.; Schmitt-Kopplin, P.; Seguin, F.; Simkus, D. N.; Shahid, A.; Takano, Y.; Thomas-Keprta, K. L.; Tripathi, H.; Weiss, G.; Zheng, Y.; Lunning, N. G.; Righter, K.; Connolly, H. C.; Lauretta, D. S. Abundant Ammonia and Nitrogen-Rich Soluble Organic Matter in Samples from Asteroid (101955) Bennu. *Nat. Astron.* **2025**, *9* (2), 199-210. https://doi.org/10.1038/s41550-024-02472-9

S4. CrysAlis PRO software; Rigaku Oxford Diffraction: 2024.

S5. Sheldrick, G. M. SHELXT – Integrated Space-Group and Crystal-Structure Determination. *Acta Crystallogr. Sect. A* **2015**, *71* (1), 3-8. https://doi.org/10.1107/S2053273314026370

S6. Sheldrick, G. M. Crystal Structure Refinement with SHELXL. *Acta Crystallogr. Sect. C* **2015**, *71* (1), 3-8. https://doi.org/10.1107/S2053229614024218

S7. Coulter, C. L. Structural Chemistry of Cyclic Nucleotides. II. Crystal and Molecular Structure of Sodium β-Cytidine 2',3'-Cyclic Phosphate. *J. Am. Chem. Soc.* **1973**, *95* (2), 570-575. https://doi.org/10.1021/ja00783a042

S8. Brandenburg, K. *Diamond*; Crystal Impact GbR: Bonn, Germany, 2022.

S9. Rout, S. K.; Wunnava, S.; Krepl, M.; Cassone, G.; Šponer, J. E.; Mast, C. B.; Powner, M. W.; Braun, D. Amino Acids Catalyse RNA Formation Under Ambient Alkaline Conditions. *Nat. Commun.* **2025**, *16* (1), 5193. https://doi.org/10.1038/s41467-025-60359-3

S10. Zgarbová, M.; Otyepka, M.; Šponer, J.; Mládek, A.; Banáš, P.; Cheatham, T. E.; Jurečka, P. Refinement of the Cornell et al. Nucleic Acids Force Field Based on Reference Quantum Chemical Calculations of Glycosidic Torsion Profiles. *J. Chem. Theory Comput.* **2011**, *7* (9), 2886-2902. https://doi.org/10.1021/ct200162x

S11. Cieplak, P.; Cornell, W. D.; Bayly, C.; Kollman, P. A. Application of the Multimolecule and Multiconformational RESP Methodology to Biopolymers: Charge Derivation for DNA,

RNA, and Proteins. *J. Comput. Chem.* **1995**, *11* (16), 1357-1377. https://doi.org/10.1002/jcc.540161106

S12. Maier, J. A.; Martinez, C.; Kasavajhala, K.; Wickstrom, L.; Hauser, K.; Simmerling, C. J. ff14SB: Improving the Accuracy of Protein Side Chain and Backbone Parameters from ff99SB. *J. Chem. Theory Comput.* **2015**, *11* (8), 3696-3713. https://doi.org/10.1021/acs.jctc.5b00255

S13. Krepl, M.; Pokorná, P.; Mlýnský, V.; Stadlbauer, P.; Šponer, J. Spontaneous Binding of Single-Stranded RNAs to RRM Proteins Visualized by Unbiased Atomistic Simulations with a Rescaled RNA Force Field. *Nucleic Acids Res.* **2022**, *50* (21), 12480–12496. https://doi.org/10.1093/nar/gkac1106

S14. Case, D. A.; Aktulga, H. M.; Belfon, K.; Ben-Shalom, I. Y.; Berryman, J. T.; Brozell, S. R.; Cerutti, D. S.; Cheatham, T. E.; Cisneros, G. A.; Cruzeiro, V. W. D.; et al. *Amber 22*; University of California: San Francisco, 2022.

S15. Krepl, M.; Vögele, J.; Kruse, H.; Duchardt-Ferner, E.; Wöhnert, J.; Šponer, J. An Intricate Balance of Hydrogen Bonding, Ion Atmosphere and Dynamics Facilitates a Seamless Uracil to Cytosine Substitution in the U-turn of the Neomycin-Sensing Riboswitch. *Nucleic Acids Res.* **2018**, *46* (13), 6528–6543. https://doi.org/10.1093/nar/gky490

S16. Ryckaert, J. P.; Ciccotti, G.; Berendsen, H. J. C. Numerical Integration of the Cartesian Equations of Motion of a System with Constraints: Molecular Dynamics of n-Alkanes. *J. Comput. Phys.* **1977**, *23* (3), 327-341. https://doi.org/10.1016/0021-9991(77)90098-5

S17. Hopkins, C. W.; Le Grand, S.; Walker, R. C.; Roitberg, A. E. Long-Time-Step Molecular Dynamics through Hydrogen Mass Repartitioning. *J. Chem. Theory Comput.* **2015**, *11* (4), 1864-1874. https://doi.org/10.1021/ct5010406

S18. Essmann, U.; Perera, L.; Berkowitz, M. L.; Darden, T.; Lee, H.; Pedersen, L. G. A Smooth Particle Mesh Ewald Method. *J. Chem. Phys.* **1995**, *103*, 8577-8593. https://doi.org/10.1063/1.470117

S19. Humphrey, W.; Dalke, A.; Schulten, K. VMD: Visual Molecular Dynamics. *J. Mol. Graph.* **1996**, *14* (1), 33-38. https://doi.org/10.1016/0263-7855(96)00018-5

S20. Roe, D. R.; Cheatham, T. E. PTRAJ and CPPTRAJ: Software for Processing and Analysis of Molecular Dynamics Trajectory Data. *J. Chem. Theory Comput.* **2013**, *9* (7), 3084-3095. https://doi.org/10.1021/ct400341p
